# Effects of chronic lithium treatment on anxious behaviors and serotonergic genes expression in the midbrain raphe nuclei in defeated male mice

**DOI:** 10.1101/2021.01.04.425168

**Authors:** Dmitry A. Smagin, Irina L. Kovalenko, Anna G. Galyamina, Irina V. Belozertseva, Nikolay V. Tamkovich, Konstantin O. Baranov, Natalia N. Kudryavtseva

## Abstract

There are experimental data that mixed anxiety/depression-like state induced by chronic social defeat stress is accompanied by development of anxiety and downregulation of serotonergic gene expression in the midbrain raphe nuclei of male mice. The paper aimed to study the effect of chronic lithium chloride (LiCl) on anxious behaviors and the expression of serotonergic genes (*Tph2*, *Slc6a4*, *Htr1a*, *Htr5b*) in the midbrain raphe nuclei of defeated mice. Slight anxiolytic effects of LiCl were found on the commucativeness in the partition test, and anxiogenic-like effects, estimated by the elevated plus-maze and social interactions tests. Chronic LiCl treatment induced overexpression of the serotonergic genes in the midbrain raphe nuclei of defeated mice. We can assume that effects of LiCl, rather anxiogenic, may be due to activation of serotonergic system induced by hyperexpression of serotonergic genes. Our findings will allow to understand the factors involved in the positive and side effects of lithium on anxiety and function of serotonergic genes which are involved into mechanisms of depression.

## Introduction

Basic and clinical studies have outlined the association between depression and neural plasticity, which is thought to be a fundamental mechanism of neuronal adaptation. Disturbances of neural plasticity (synaptic plasticity, neurogenesis, energetic metabolism, *etc.*) play a significant role in the development of depression [1, 2, 3]. The central role of the brain serotonergic system in the mechanisms of stress, anxiety, depression and bipolar disorder [4–6], on the one hand, and in the neural plasticity [2, 3, 7, 8], on the other hand, was shown in numerous studies. It is widely believed that serotonergic dysbalance is a key pathophysiological mechanism in major depression. Moreover, deficiency in serotonin synaptic signaling fundamentally impacts the pathophysiology of different disorders [9].

Our experiments revealed that chronic social defeat stress (CSDS) in daily agonistic interactions of C57BL/6 male mice leads to the development of the mixed anxiety/depression-like state, which is similar to as judged by similarities in symptoms, etiology, sensitivity to antidepressants and anxiolytics as well as neurochemical changes in the depressive patients [10, 11, 12]. It has been shown that at the stage of pronounced depression-like state (20 days of CSDS) a decreased brain serotonergic function at the level of synthesis, metabolism and receptor sensitivity were found in the defeated mice. Hypofunction and, possibly, depletion of the brain serotonergic system was assumed to be a result of its prolonged activation under CSDS in animals [12]. This hypofunction of the serotonergic system is accompanied by downregulation of expression of serotonergic *Tph2, Slc6a4, Maoa* and *Htr1a* genes [13] encoding the TPH, the rate limiting enzyme of the serotonin (5-HT) pathway, serotonin transporter, monoamine oxidase A (MAOA), and 5HT_1A_ receptors, respectively, in the midbrain raphe nuclei, containing of perycaryons of serotonergic neurons. After a period of relative rest without agonistic interactions serotonergic genes continued to be expressed at reduced rates for at least two weeks [13]. Later, whole transcriptome analysis (RNA-seq) in similar experiments confirmed that the *Tph2, Ddc, Htr2a*, *Htr3a, Htr5b* genes were expressed at lower levels in depressed mice [14]. Our findings allowed to suggest that the low efficacy of traditional treatment by antidepressants, primarily by inhibitors of serotonin transporter reuptake, as well as frequent relapses of the disease after a seemingly successful treatment can be explained by altered expression of the genes, in particular serotonergic ones, which remain downregulated. Obviously, it is necessary to look for pharmacological agents that can accommodate restoring the gene expression altered in the course of disease development.

To choose a medicine as a likely candidate capable of restoring gene expression we considered lithium, which acts through a totally different mechanism. Besides, lithium salts are widely used in psychiatric practice in monotherapy regimens as mood stabilizers [15–18], in the supportive therapy of psychoemotional disorders [19], in prevention suicidal behavior in patients with manic-depressive psychosis [20–22]. Lithium is also used at onset of depressive phase in patients with bipolar disorder, for prevention of mood disorders [15, 23–26], or relapses in schizophrenia patients with aggressive or suicidal behavior, convulsions *etc* [27–30].

There is evidence that therapeutic action of lithium is due to the effects on serotonergic neurotransmission [4, 31]. The studies on humans demonstrated that the effects of lithium on the serotonergic system depend on tryptophan hydroxylase protein (TPH) variants [32], and that lithium may act through 5-HT_1B_ receptors as shown in an animal models [33]. Regular administration of lithium increases the density of the serotonin uptake site in cortical regions suggesting an increase in the number of serotonin transporters in the brain regions containing nerve terminals of serotonergic neurons [34]. In our study the behavioral effects of lithium-based enterosorbent, called ‘Noolit’, on male mice with mixed depression/anxiety-like state produced obvious anxiolytic and antidepressant effects [35].

In this study, we took into account a lot of recently obtained data on the existence of lithium-sensitive genes that may be involved in the development of affective and neurodegenerative disorders [36–42]. The aim of this work was to study the effect of chronic administration of lithium chloride on anxiety behavior and expression of serotonergic genes (*Tph2, Slc6a4, Htr1a, Htr5b*) in the midbrain nuclei of defeated male mice. Since lithium is used at the initial stage of the depressive phase of bipolar disorder and for the prevention of mood diseases in patients [15, 23–26], in our experiment we administered lithium during the period of repeated agonistic interactions, expecting to see its protective effects shown earlier [43–44].

## Materials and Methods

### Animals

Adult C57BL/6 male mice were obtained from Animal Breeding Facility, Branch of Institute of Bioorganic Chemistry of the RAS (Pushchino, Moscow region, Russia). The animals were housed under standard conditions (12:12 h light/dark regime, switch-on at 8.00 a.m.; food (pellets) and water available *ad libitum*). Mice were weaned at one month of age and housed in groups of 8-10 in plastic cages. All procedures were carried out in compliance with international regulations for animal experiments (Directive 2010/63/EU of the European Parliament and of the Council on the Protection of Animals Used for Scientific Purposes). The protocol for the studies was approved by Scientific Council No 9 of the Institute of Cytology and Genetics SD RAS of March, 24, 2010, N 613 (Novosibirsk).

### Behavioral study

Prolonged experience of chronic social defeat stress, accompanied by strong anxiety in male mice was induced using the sensory contact model [10, 44]. Pairs of weight-matched animals were each placed in a cage (28 ×14 ×10 cm) bisected by a perforated transparent partition allowing the animals to see, hear and smell each other, but preventing physical contact. The animals were left undisturbed for two or three days to adapt to new housing conditions and sensory contact before they were exposed to encounter. Every afternoon (14:00-17:00 p.m. local time), the cage lid was replaced by a transparent one, and 5 min later (the period necessary for mouse activation), the partition was removed for 10 minutes to encourage agonistic interactions. The superiority of one of the mice was firmly established within two or three encounters with the same opponent. The superior mouse would be attacking, biting and chasing another, who would be displaying only defensive behavior (upright postures, sideways postures, freezing or withdrawal, lying on the back). As a rule, agonistic interactions between males are discontinued by lowering the partition if the strong aggression has lasted 3 min, in some cases, less. Each defeated mouse (defeater, loser) was exposed to the same aggressive mice for three days, while afterwards each loser was placed, once a day after the fight, in an unfamiliar cage with an unfamiliar winner behind the partition. Each winning mouse (winners, aggressors) remained in its original cage. This procedure was performed once a day for 20 days and yielded an equal number of winners and losers.

After 6 days of social stress, defeated mice were treated with saline or lithium chloride (LiCl; Merck, Germany) at a dose of 100 mg/kg (in behavioral study), i.p., once a day in the morning at 9-10 AM, 16 days. Agonistic interactions were performed in afternoon including days of behavioral testing.

In the experiment for the study of the effect of chronic lithium treatment on serotonergic gene expression in the midbrain raphe nuclei, we used a group of animals after treatment with LiCl (150 mg/kg, two weeks daily) that did not undergo behavioral testing to avoid stress effects. In an additional experiment, we found that both doses of LiCl (100 and 150 mg/kg) had the same effect on the behavior of the animals in the elevated plus-maze test. The control mice and losers the day after the last agonistic interactions, on the 21st day, were decapitated.

Three groups of animals were used in the experiment: the controls — mice without consecutive experiences of social defeats; defeated males after chronic treatment with saline (Sal-treated losers); defeated males after chronic treatment with LiCl (LiCl-treated losers). The intact state of animals, which is used in our pharmacological experiments, makes it possible to study the effectiveness of the preparations. After two weeks of treatment on the background of continuing agonistic interactions the behavior of animals was evaluated (one test/day during three days, Fig. 1) in the partition, elevated plus-maze, exploratory activity and social interactions tests, which were used for measuring the level of anxiety in different experimental situations.

**Figure 1.**
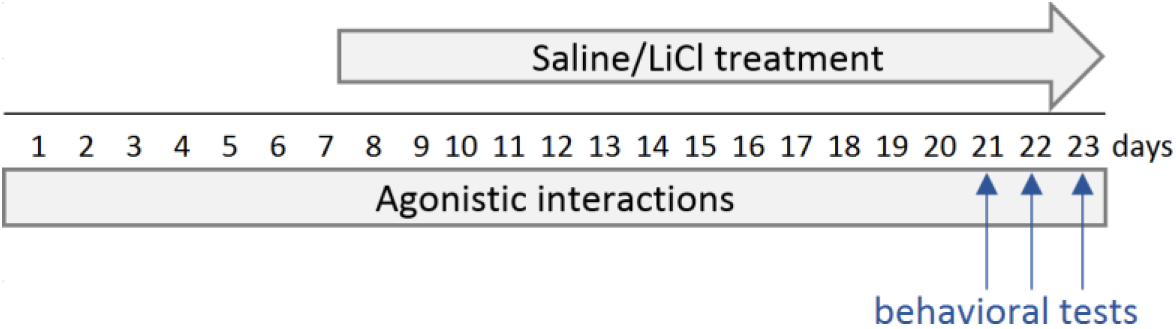
Scheme of experiment: chronic LiCl and/or saline treatment of defeated mice. Behavioral tests: Day 21 – the partition test and agonistic interaction; Day 22 – the elevated plus-maze test, Day 23 – the social interaction test.

Two different groups (the controls, saline- and LiCl –treated losers) were used: 1) for the study of effect the LiCl on anxious behavior in different behavioral tests; 2) for molecular study of gene expression in the brain in order to avoid the effect of additional non-specific influences of in behavioral tests, for example, stress.

### Behavioral tests

#### The partition test

Partition can be utilized as a tool for estimation of mouse behavioral reaction to a conspecific behind the transparent perforated partition dividing the experimental cage into the equal parts [45]. The number of approaches to the partition, and the total time spent near it (moving near the partition, smelling and touching it with nose, one or two paws, putting noses into the holes) are scored during 5 min as indices of reacting to the partner. The time the males show sideways position or “turning away” near the partition is not included in the total time of test. The experimental procedure is as follows: the winners and losers live together in a cage with a partition. On the testing day, the lid of the cage is replaced by a transparent one. 5 minutes later (period of activation) behavioral responses of the losers and the controls toward the familiar partner is recorded for 5 min. This test is used for the study of communicativeness (sociability) as well as level of anxiety: it was shown that decrease of partition items correlates positively with indices of anxiety estimated in plus-maze test [46].

#### Elevated plus-maze test [46]

The elevated plus-maze consisted of two open arms (25× 5 cm) and two closed arms (25×5×15 cm) and was placed in a dimly lit room. The two arms of each type were opposite to each other and extended from a central platform (5×5 cm). The maze was elevated to a 50 cm above floor.

The cover of the experimental cage with a mouse was replaced by transparent lid in the same room 5 min before exposure to the plus-maze. The mouse was placed with the nose to the closed arm at the central platform. The following measures were recorded during 5-min of plus-maze test: 1) total entries; 2) open arm entries (four paws in open arm), closed arm entries (four paws in closed arm) and central platform entries; 3) time spent in open arms, closed arms, and central platform (center); 4) the number of passages from one closed arm to another; 5) the number of head-dips (looking down on the floor below the plus-maze); 6) the number of peeping when mouse being in closed arms (the head of the mouse is put out from the closed arm and returns quickly back). Indices 2 and 3 are considered as measures of the level of anxiety, indices 1 and 4 are related to locomotor activity, indices 5 and 6 are considered as risk assessment behavior. Time spent in the closed, open arms and in the central platform (center) was calculated in percentages from total testing time. Plus-maze was thoroughly cleaned between sessions. This test we consider as low stressful test (test with low aversion).

#### Exploratory Activity and Social Interaction tests [47]

The open-field (36×23 cm) with perforated turned metal wire holder (inverted pencil holder, bottom diameter - 10.5 cm) in one of the cage corners was used. Each mouse was placed individually in the opposite from holder corner of the open field for 5 min. This test allows to estimate the exploratory behavior of mice in novel conditions with unfamiliar object - wire holder (exploratory activity test). In comparison with plus-maze test with low aversion in dimly lit room, this test can be considered as higher stressful situation. Then an unfamiliar group housed male was placed under the wire holder for 5 min to study the reaction of male mice to conspecific in familiar situation (Social interaction test). The following behavioral variables were registered:

1. Automatic registration with EthoVision XT software (version 11.0; Noldus Information Technology, The Netherlands) of the tracking score (distance) during testing time with differentiation of place near the wire holder (5 cm around) of the cage and total time spent in the opposite to wire holder corner;
2. Manual registration with Observer XT (version 7.0; Noldus Information Technology, The Netherlands) was used for following behavior indicators of communicativeness/sociability: 1) the number and/or duration of 1) rearing (exploratory activity); 2) grooming (self-oriented behavior — licking of the fur on the flanks or abdomen, washing over the head from ear to snout); 3) approaches to the wire holder and the total time (s) spent near it (moving near the wire holder, smelling and touching it with nose, one or more paws, getting on a wire holder and a contact with it by four paws). The duration of sideways position or “turning away” near the wire holder was not included in the total time. After each test, the open field and wire holder apparatus were thoroughly washed and dried with napkins.

Preliminary analysis of LiCl-treated losers’ behavior in exploratory activity test clearly divided animals into two groups — the LiCl+ sensitive (LiCl+) and LiCl-less sensitive (LiCl-) losers to the chronic LiCl treatment. The LiCl+treated anxious losers, when they were placed in an unfamiliar cage, seated in the corner only and did not explore the cage. As behavioral parameter for division into groups was taken avoidance of novel object (wire holder): LiCl+ — mice did not approach to wire holder at all in comparison with other LiCl-treated animals. In the control and Sal-treated losers, we observed a natural exploratory activity in novel conditions.

#### RT PCR method

To measure mRNA levels of serotonergic genes in the midbrain raphe nuclei, all the mice were studied: 21-day losers after chronic LiCl or saline injections, 24 hours after the last agonistic interaction, and the controls. The midbrain raphe nuclei area was dissected according to the Mouse Brain Atlas [http://mouse.brain-map.org/static/atlas] in animals of all experimental groups. The brain regions were removed and chilled rapidly on ice. Dissection of the brain regions was made by the same experimenter. All biological samples were encrypted, were rapidly frozen in liquid nitrogen and stored at −70° C until use.

The determination of serotonin gene expression was done at Biolabmix (Novosibirsk, RF). Total RNA was isolated from the tissues using TRIzol (Invitrogen) according to the manufacturer’s instructions. The concentration of total RNA was quantified by measuring the absorbance at 260 nm. The integrity of total RNA was assessed using agarose gel electrophoresis. cDNAs were synthesized using total RNA (1 μg) and M-MuLV –RH First Strand cDNA Synthesis Kit (Biolabmix) random N9 primer (100 ng) and MoMLV reverse transcriptase (200 U, Biosan). Each RT reaction was run in duplicate. RNA aliquots were used to confirm the absence of any genomic DNA in each sample.

The *Tph2, Htr1a, Htr5b, Slc6a4* and *B2M* cDNA levels were quantified by SybrGreen-based real-time PCR in a total volume of 25 μl containing an aliquot of the RT mixture, master mix Biomaster HS-qPCR SYBR Blue (2×) (Biolabmix, Novosibirsk, RF) and specific primers (300 nM). dNTPs (200 nM), F and R primers (300 nM), SybrGreen I (1:20000, Invitrogen), standard PCR buffer, and hot-start TaqDNA polymerase (0.5 U, Biosan). Amplification was run for 3 min at 95°C followed by 40 cycles of 6 s at 96°C, 6 s at 60°C, 12 s at 72°C. Fluorescence was monitored for 5 s after each cycle at the appropriate melting temperature. To check for the presence of non-specific PCR products or primer dimer, a melting curve analysis was performed after the final PCR cycle.

Amplification efficiencies were calculated using a relative standard curve derived from threefold serial dilutions of pooled cDNA. In all cases, the amplification efficiency was higher than 90%. Each sample was PCR-amplified twice. qRT-PCR results were quantified using the relative standard curve method. Samples from animals of different experimental groups and their brain areas were mixed before molecular study. Extraction of total RNA, reverse transcription and qRT-PCR was made during 2 weeks by one experimenter in blind regimen with use of one buffer and primers set.

#### Statistical analysis

Statistical analysis for behavioral data was performed using either one-way ANOVA for parametric variables or Kruskal-Wallis test with factor “group” (three levels: the control group, Sal-treated losers, LiCl-treated losers) followed by the Tukey’s multiple comparisons post hoc test for parametric variables or Kruskal-Wallis test with Dunn’s multiple comparisons post hoc if the parametric assumptions were not satisfied. To display the variability among the values, the data are presented as a box-whisker plot showing means (*plus sign*), medians (*solid lines*) and 25%/75% quartiles with whiskers indicating 10th and 90th percentiles. All statistical analyses were performed using XLStat software (Addinsoft, www.xlstat.com). For each experimental sample, the relative amount of mRNA is determined from the appropriate standard curve. The *B2m* gene expression has been used to normalize the expression of serotonergic genes. Measurement statistics were provided by BioRad Amplifier software (USA).

## Results

### Effects of chronic LiCl treatment on the behavior of defeated mice in the partition test

(Fig. 2) One-way ANOVA revealed the influence of the “group” factor on the number of approaches to the partition as reaction to the partner in the neighboring compartment of cage (F (2, 33) = 6.619, *P* = 0.0038) and rearing (F (2, 33) = 3.293, *P* = 0.0496). Kruskal-Wallis test revealed the influence of the “group” factor on the total time spent near the partition (H = 9.816, *P* = 0.0074). Tukey’s multiple comparisons test was used for the number of approaches and the number of rearing and Dunn’s multiple comparisons test for the time spent near the partition. For the Sal-treated losers all parameters were significantly lower in comparison with the controls for the number of approaches (*P* = 0.0027), total time spent near the partition (*P* = 0.0053) and number of rearing (*P* = 0.0390).

**Figure 2.**
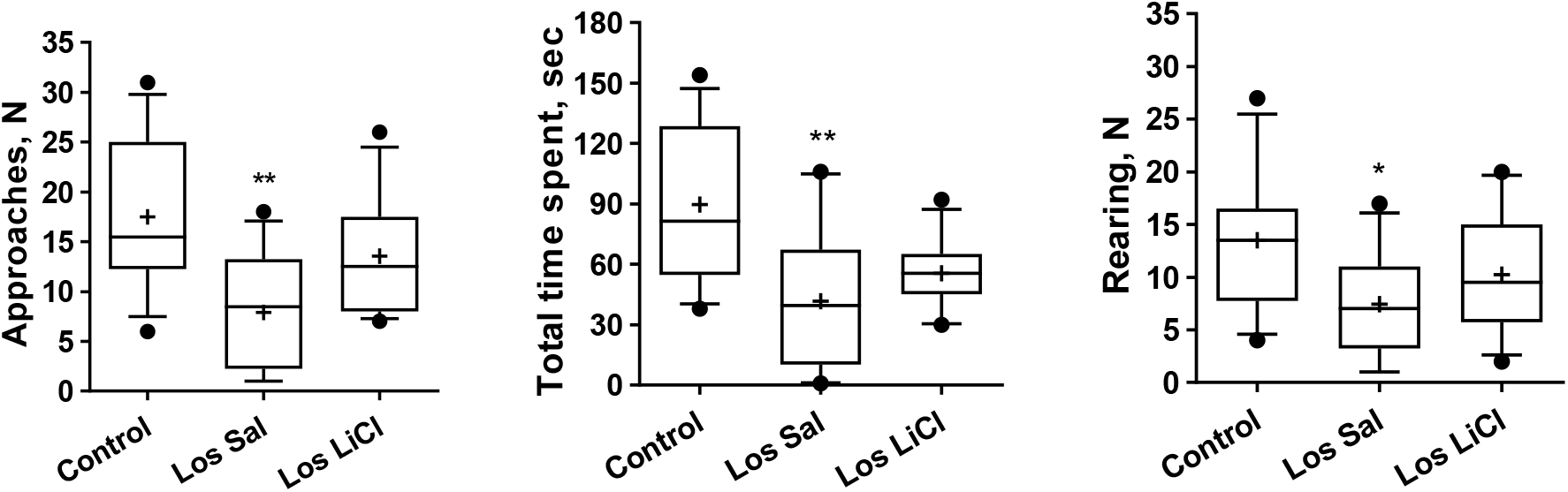
Effects of chronic LiCl treatment on the behavior of defeated mice in the partition test. Los Sal — Sal-treated losers, Los LiCl — LiCl-treated losers. Values are shown as means (*plus sign*) and medians (*solid lines*) and 25%/75% quartiles in a box-whisker plot with whiskers indicating 10th and 90th percentiles. * *P* < 0.05, ** *P* < 0.01 *vs* the controls; N=12 for each group.

Thus, Sal-treated losers, as expected, had lower communicativeness toward the partner in the neighboring compartment, as revealed number to and total time spent near the partition, and decreased exploratory activity estimated by the rearing behavior. Under the LiCl treatment, all parameters did not yet differ significantly from the control. We can assume that LiCl induced slight anxiolytic effects in the losers.

### Effects of chronic LiCl treatment on the behavior of defeated mice in the plus-maze test

One-way ANOVA revealed a significant influence of the factor “group” (the control, Sal-treated losers, LiCl-treated losers) on the number of central platform entries (F (2, 31) = 4.130, *P* = 0.0257) and number of closed arm entries (F (2, 31) = 4.661, *P* = 0.0170) as well as on the number of passages (F (2, 31) = 4.717, *P* = 0.0163), head dips (F (2, 31) = 3.590, *P* = 0.0396), total entries (F (2, 31) = 4.552, *P* = 0.0185). Based on the Tukey’s multiple comparisons post hoc test revealed in the LiCl-treated group, that number of central platform entries (*P* = 0.0277) and closed arm entries (*P* = 0.0185), number of passages (*P* = 0.0166), head dips (*P* = 0.0403), and total entries (*P* = 0.0192) were lower in comparisom with the respective behavioral items in the controls (Fig. 3).

**Figure 3.**
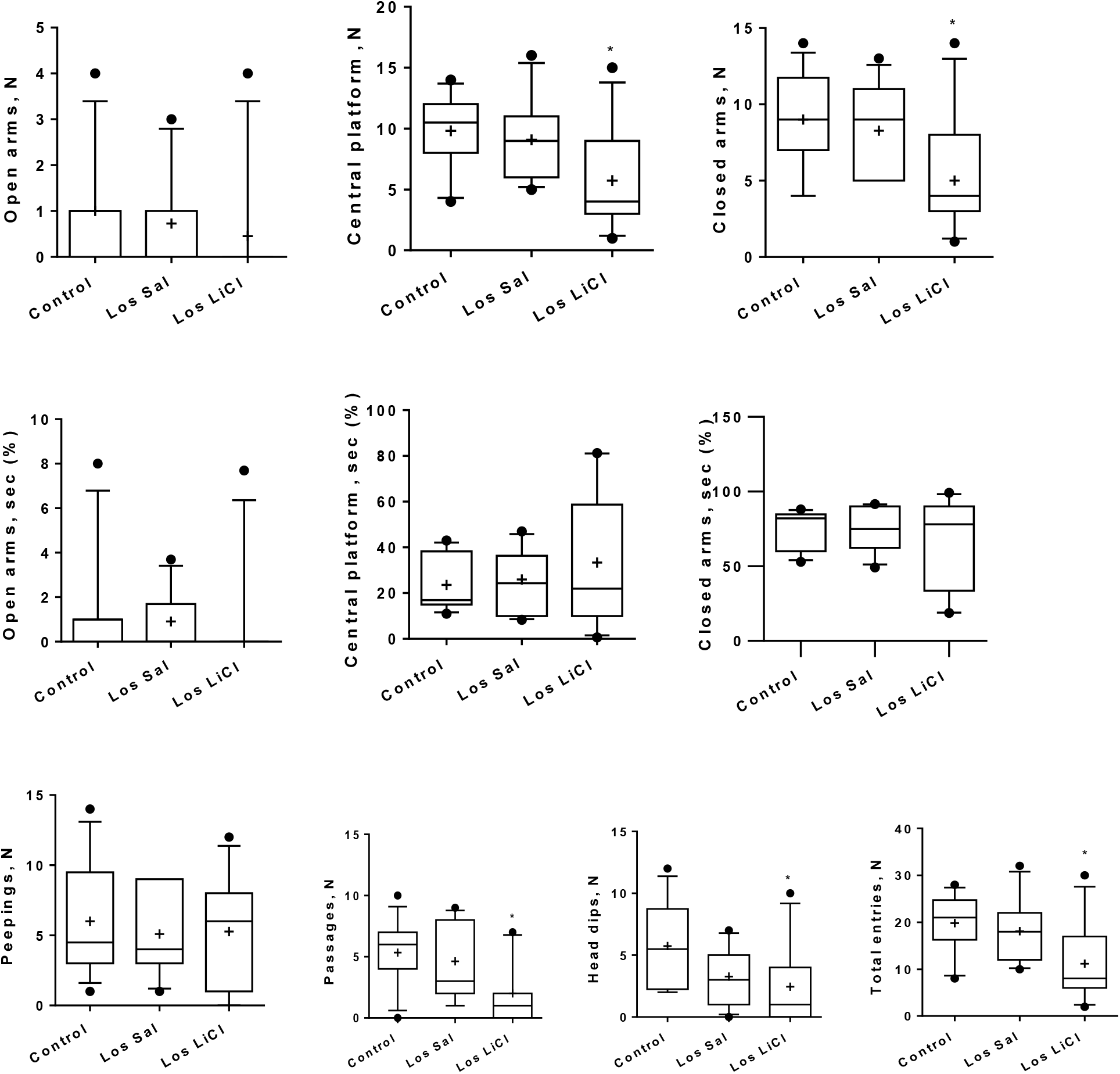
Effects of chronic LiCl treatment on the behavior of defeated mice in the plus-maze test. Los Sal – Sal-treated losers, Los LiCl – LiCl-treated losers. * — *P* < 0.05 *vs* controls, Tukey’s multiple comparisons post hoc test; N = 12 for each group. Values are shown as means (*plus sign*) and medians (*solid lines*) and 25%/75% quartiles in box-whisker plot with whiskers indicating 10th and 90th percentile. * *P* < 0.05 *vs* the control group (*Tukey’s multiple comparisons post hoc test*).

Previously, it was repeatedly shown that expressed anxiety is developed in the losers after 20-day social defeat stress, estimated by the elevated plus-maze test [12, 43]. Against the background of the chronic saline treatment, these differences were less pronounced. A cautious conclusion can be made that the saline *per se* has a protective antitoxic effect under chronic injections. However, we can assume, that the decrease of locomotor activity under chronic LiCl treatment (decreased number of total entries, passages and head-dips) may be considered as anxiogenic effects, since behavioral deficit, fear and indifference are considered as main symptoms of mixed anxiety/depression-like state.

### Effects of chronic LiCl treatment on the exploratory activity of the defeated mice with different sensitivity to LiCl treatment in novel situation toward novel object

(empty wire holder, Fig. 4) According to the level of wire holder avoidance (see description in Materials and Methods) we divided LiCl-treated losers into two subgroups: LiCl+sensitive and LiCl-sensitive losers (less sensitive). One-way ANOVA revealed a significant influence of the factor “group” (the control, Sal-treated, LiCl-treated, LiCl+treated losers) on the total tracking (F (3, 29) = 26.50, *P* < 0.0001) (tracking, cm). Tukey’s multiple comparisons test revealed the difference between the following groups: Control *vs* Sal-treated (*P* = 0.0003), LiCl-treated (*P* < 0.0001) and LiCl+treated losers (*P* < 0.0001); LiCl+treated *vs* Sal-treated losers (*P* = 0.0013).

**Figure 4.**
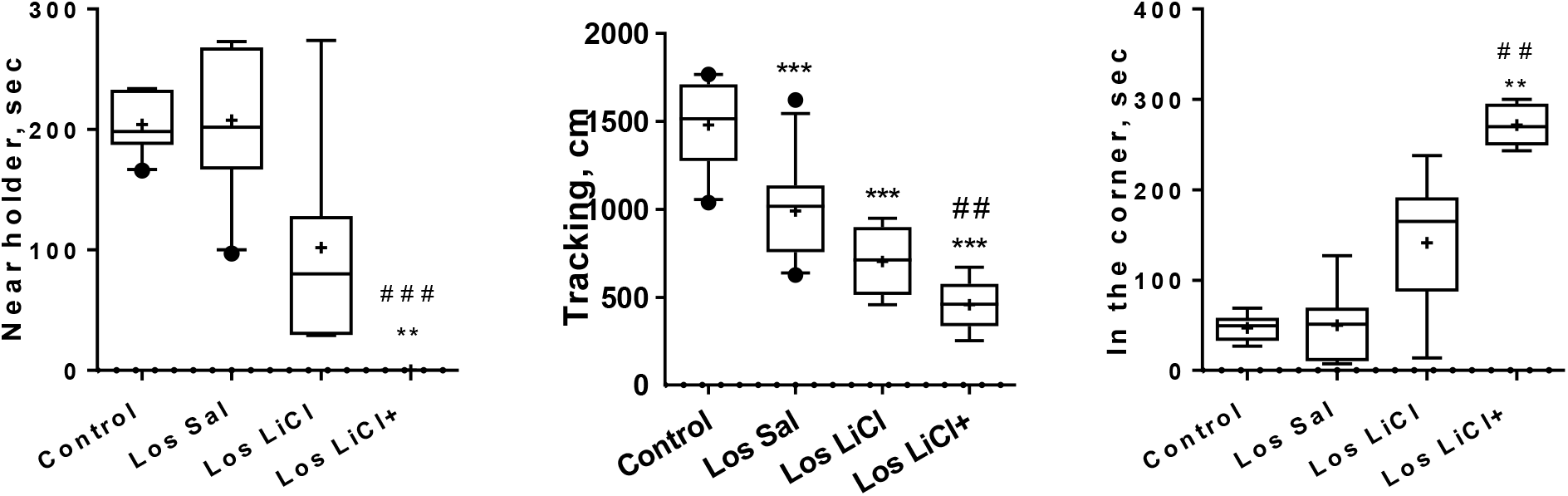
Effects of LiCl on the exploratory behavior of defeated mice in new situation and toward novel object (holder). Los Sal – Sal-treated losers, Los LiCl – LiCl-treated losers. Values are shown as means (*plus sign*) and medians (*solid lines*) and 25%/75% quartiles in box-whisker plot with whiskers indicating 10th and 90th percentile. ** — *P* < 0.01, *** — *P* < 0.01 *vs* controls; ## — *P* < 0.01, ### - *P* < 0.001 *vs* Sal-treated losers (*Tukey’s multiple comparisons post hoc test*); Control (N=9); Sal-treated losers (N=11), LiCl-treated losers (N=7), LiCl+treated losers (N=5).

Kruskal-Wallis test revealed the influence of the “group” factor on the time spent *in the corner* (H = 18.03, *P* = 0.0004) and near *the wire holder* (H = 17.56, *P* = 0.0005). Dunn’s multiple comparisons test revealed the difference between the following parameters (Fig. 4): *in the corner* (sec) Control *vs* LiCl+treated losers (*P* = 0.0027) and Sal-treated losers *vs* LiCl+treated losers (*P* = 0.0012); *near the wire holder* (sec) Control *vs* LiCl+treated losers (*P* = 0.0040) and Sal-treated losers *vs* LiCl+treated losers (*P* = 0.0010).

It has been shown that LiCl+treated losers were more sensitive to effects of LiCl and displayed the less exploratory activity, estimated by total tracking time. Major time they spent in the corner and never were near wire holder. Together with behavior in the plus-maze and partition tests these data may indicate decreased exploratory activity and increased level of anxiety under chronic LiCl treatment.

### Effect chronic LiCl treatment on the reaction of mice to unfamiliar partner in the Social interaction test (Fig. 5)

This test we considered as measuring the sociability with the unfamiliar partner in conditions which has already become familiar during 5 minutes before introduction of partner. During previous 5 minutes the mice realized that they were not in danger, and they began slightly to examine the cage.

**Figure 5.**
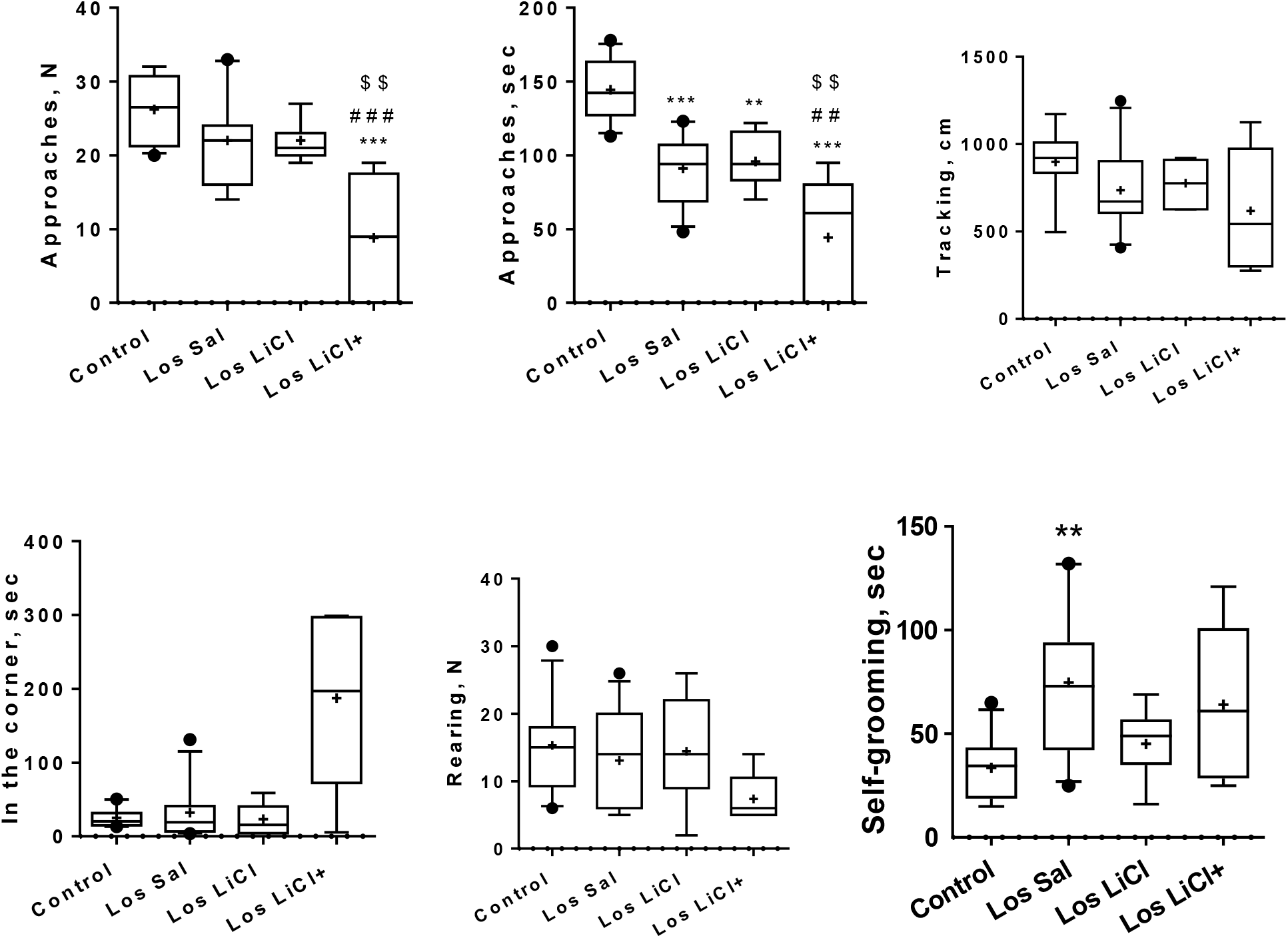
Effects of LiCl on the behavior of defeated mice in the social interaction test as a reaction to the partner under wire holder. Los Sal – Sal-treated losers, Los LiCl – LiCl-treated losers. Values are shown as means (*plus sign*) and medians (*solid lines*) and 25%/75% quartiles in box-whisker plot with whiskers indicating 10th and 90th percentile; For the control – N=9, Sal-treated losers – N=11, LiCl-treated losers – N=7), LiCl+treated losers – N=5. ** — *P* < 0.01, *** — *P* < 0.01 *vs* controls; ## — *P* < 0.01, ### — *P* < 0.001 *vs* Sal-treated losers; $$ — *P* < 0.01 *vs* LiCl-treated losers.

One-way ANOVA indicate a significant influence of the factor “group” (Control, Sal-treated, LiCl-treated, LiCl+treated losers) on the number of *approaches* to partner (F (3, 31) = 11.35, *P* < 0.0001) and *total time spent* near the wire holder (Approaches, sec) (F (3, 31) = 20.98, *P* < 0.0001). Kruskal-Wallis test revealed the influence of the “group” factor on the time of *self-grooming* (H = 9.816, *P* = 0.0212).

Tukey’s multiple comparisons test was used for the following behavioral parameters (Figure 5): *approaches* (N) - Control *vs* LiCl+treated losers (*P* < 0.0001), Sal-treated loser *vs* LiCl+ treated losers (*P* = 0.0008), LiCl-treated *vs* LiCl+ treated losers (*P* = 0,0020); *approaches* (sec) — the control *vs* Sal-treated losers (*P* < 0.0001), LiCl-treated losers (*P* = 0.0017), LiCl+ treated losers (*P* < 0.0001); Sal-treated losers *vs* LiCl+ treated losers (*P* = 0.0083), LiCl-treated losers *vs* LiCl+ treated losers (*P* = 0,0073). Dunn’s multiple comparisons test was used for the time of *self-grooming* (sec) — the control *vs* Sal-treated losers (*P* < 0.0133).

Chronic LiCl treatment induced strong anxiogenic effect in LiCl+treated losers in comparison with the control and Sal-treated losers which was estimated by decreased number of approaches to and time spent near the holder with partner (approaches, sec). Time spent in the corners was significantly higher in comparison with all groups. Anxiogenic effects in LiCl-treated losers was significantly less differing in comparison with the control mice.

### Effects of chronic LiCl treatment on the expression of serotonergic genes in the midbrain raphe nuclei of defeated mice (Fig. 6)

The measurement data is provided by the BioRad Amplifier software (USA). Chronic LiCl treatment induced overexpression of the *Tph2* gene, encoded the limiting enzyme of serotonin synthesis, in comparison with the controls and Sal-treated losers (both *P* < 0.01); the *Slc6a4* gene, encoded serotonin transporter in comparison with the control and Sal-treated losers (*P* < 0.05 and *P* < 0.01, respectively); the *Htr1a* gene, encoded 5HT_1a_ receptors, in comparison with the controls and Sal-treated losers (for both *P* < 0.01); and the *Htr5b* genes, encoded 5HT_5b_ receptors, in comparison with the controls and Sal-treated losers (*P* < 0.01 and *P* < 0.05, respectively) in the midbrain raphe nuclei of defeated mice.

**Figure 6.**
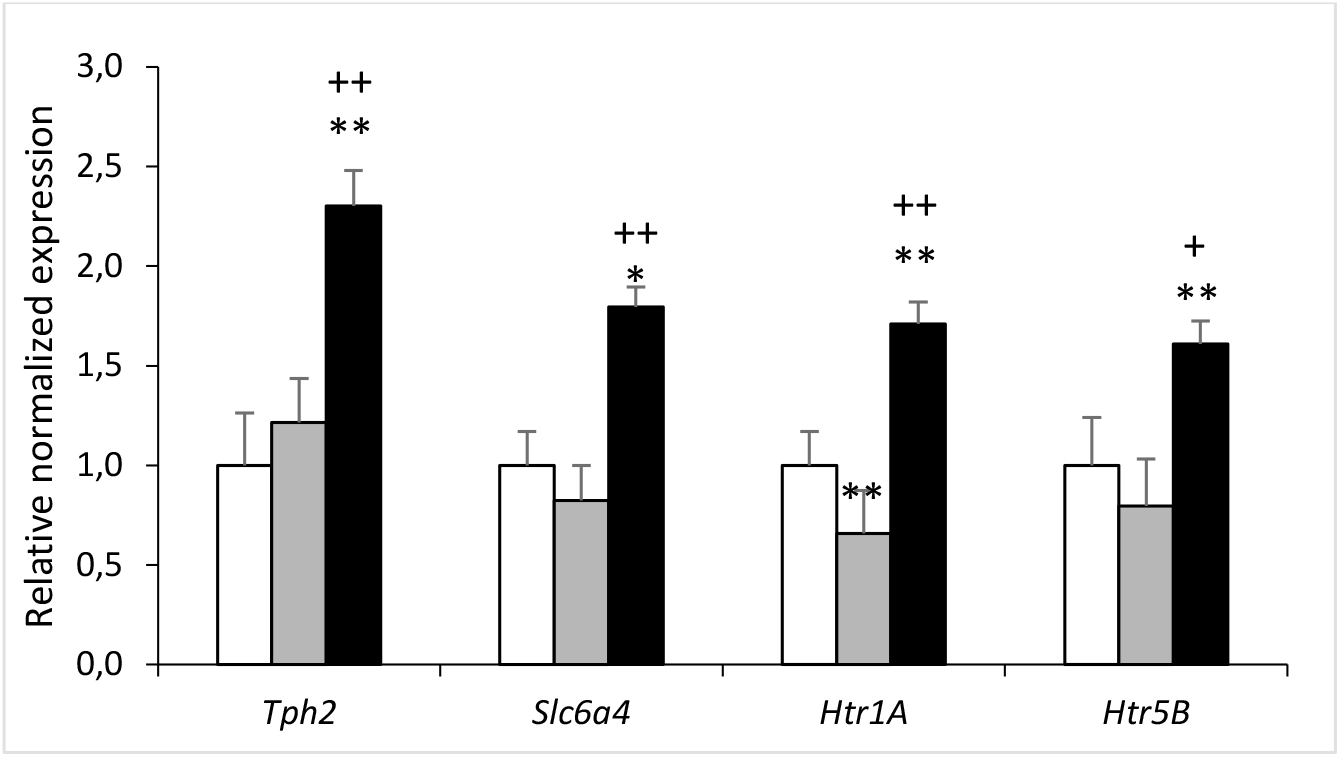
Influence of chronic LiCl treatment on serotonergic gene expression in the midbrain raphe nuclei of defeated mice. The measurement data is provided by the BioRad Amplifier software (USA). White columns – the control (N=9); grey columns – Sal-treated losers (N = 7); black columns – LiCl-treated losers (N=9); * – *P* < 0.05, ** – *P* < 0.01 *vs* controls; + – *P* < 0.05, ++ – *P* < 0.01*vs* Sal-treated losers, t-Student test. The data are reported as mean ± SEM.

Some discrepancy between the data obtained in (Boyarskikh et al., 2013) and the data of these studies can be explained by the lack of complete identity of these groups. In 2013, we did not administer saline to a group of the losers. In this experiment, saline was administered daily for two weeks. There is also the possibility that saline itself may have a protective antitoxic effect with chronic injections decreasing effect of CSDS.

## Discussion

The discussion is focused on three aspects: a) the causes of varying sensitivity of animals to the effects of pharmacological agents, in particular, LiCl; b) the effects of LiCl on psychoemotional states of animals and humans; c) varying effects of LiCl treatment in dependence on brain serotonergic activity in the mice with different social experiences in agonistic interactions.

### a. Causes of varying sensitivity of animals to the effects of pharmacological agents

A very important aspect is the varied sensitivity of animals of the same group to drugs, in particular in this work, to LiCl. When studying the behavior of aggressive [48] and defeated male mice (this experiment) in social interaction test we found that animals of group split into subgroups according to different sensitivity to LiCl. Naturally, the question arises: why inbred animals split into sensitive and less sensitive groups to drug effect under obviously identical experimental conditions? One of the plausible reasons, as supposed in [49–50], could be the differences in prenatal and early postnatal development, which have been overlooked in the standardized experimental setting.

However, in our opinion, a more possible assumption is that the baseline psychoemotional statuses of experimental animals grown in group-housed conditions in a vivarium are different. Mice are known to form despotic dominance hierarchy with one male being dominant and others being subordinates [51]. Social status leaves an imprint on the behavior and brain neurochemistry of mice. In this study we got additional evidence that the effect of a drug may depend on the psychoemotional state of an individual. In other words, the neurochemical background can modify the effect of a drug, sometimes to the opposite of the expected effect.

Moreover, some effects of a drug can be detected in one situation (test) and not manifested in another. Apparently the cause is that the leading motivation developed in the experimental conditions underlies many (but not all) forms of behavior. Sometimes it could be a struggle between two opposite motivations (ambivalence), for example, fear and communicativeness (the test of social interactions), or anxiety and exploratory activity in a new situation (the plus-maze test). The balance between the leading motivational components is situational, is expressed through a psychoemotional state, and, consequently, the underlying neurochemical background that may mediate the effect of the drug. The division of male mice into the subgroups, susceptible or resilient to the effects of CSDS, was also observed by other researchers [52–54 *etc*.].

### b. Effects of LiCl on psychoemotional state of animals and humans

In our behavioral studies chronic injections of intact male mice with LiCl for two weeks was shown to produce anxiolytic effects: the anxiety level estimated in the plus-maze and social interaction tests decreased [48]. In the present experiment, under preventive treatment of defeated male mice on the background of severe anxiety, the effect of LiCl correlated with the behavioral scores in different tests. Slight anxiolytic effects were observed in the home cage in the partition test: interest to an unfamiliar partner (sociability) in the neighboring compartment, estimated by the number of approaches to and total time spent near the partition was decreased in Sal-treated losers in comparison with the control, but did not differ from the control in mice after LiCl treatment. The LiCl produced decrease in the number of passages, total entries and head-dips, as well as in the number of central platform and closed arms entries which can readily be explained in the context of general behavioral deficit characteristic of mixed anxiety/depression-like state, which develops in animals under CSDS [10–12]. This effect of LiCl may be considered as pro-depressive effects in this experimental context.

Apparent anxiogenic effects on LiCl-treated losers in the social interaction test may be a result of a frightening situation. In the group of defeated mice LiCl produced marked variable effects: about 40% of the LiCl+treated losers demonstrated pronounced anxiety behavior in new situation toward novel object and unfamiliar partner under wire holder. They spent much time in the corner opposite to the wire holder corner, and never near it. Total tracking time was also decreased in all groups of the losers in comparison with the control with most decreased parameter in the LiCl+sensitive losers. An unfamiliar partner under the wire holder induced no interest in the male mice but rather fear: number and total time of approaches were significantly less in comparison with other experimental groups.

Earlier in similar experiments, it was shown that repeated aggression experiences is accompanied by the development of a whole range of changes in behavior and psychoemotional state with symptoms of psychosis-like behavior: the aggressive males demonstrated an increased number of stereotypical behaviors (self-grooming, rotation, jerks etc), development of anxiety, hyperactivity and strong aggressive motivation, which, along with other signs, indicated the development of a pathological behavior [44, 55]. The LiCl was administered preventively to aggressive males on the background of agonistic interactions, as well as therapeutically to males with the 20-day period of repeated aggression during the period without agonistic interactions [48].

Preventive chronic LiCl injections to aggressive male mice were shown to induce pronounced anxiogenic effects similar to those in the losers: LiCl further enhanced anxiety, which was shown earlier for male mice with repeated experience of aggression in the partition test and elevated plus-maze test [48, 56]. In the social interaction test about 40% of aggressive mice demonstrated pronounced anxiety following LiCl treatment with a decrease in communicativeness and exploratory activity as compared with the controls. Therefore, the anxiogenic effect is likely to be consequence of the stress that accompanies agonistic interactions.

Under therapeutic treatment of aggressive males with the pathology formed during the fight period, the anxiolytic effect of LiCl became evident when analyzing the findings of the social interaction test, namely the reaction to a partner, which was characterized bу an increased interest to the partner under the wire holder [48]. However, in the plus-maze and partition tests no effects of LiCl were detected. Thus, we concluded that chronic LiCl treatment could produce opposite effects in intact animals and under preventive and therapeutic treatment of males with repeated experience of aggression. There were no effects on the expression of aggressive behavior.

In the study of the diazepam effect, similar data were obtained in slightly different experimental context: acute administration of the drug to mice with a short-term experience of aggression induced anxiogenic effect, whereas in the males with a long-term experience of aggression diazepam had an anxiolytic effect [57]. Similar data were obtained in other work: the effects of the anxiolytic chlordiazepoxide on aggressive behavior of animals with different social status may differ [58].

Thus, the effects of chronic LiCl treatment can depend on the method of treatment (preventive, therapeutic), psychoemotional state developing under positive or negative social experience of animals (intact, aggressive, defeated), type of behavior (tests), and can have both anxiogenic and anxiolytic effect or no effect whatsoever as shown in our study (Table 1).

**Table 1.**
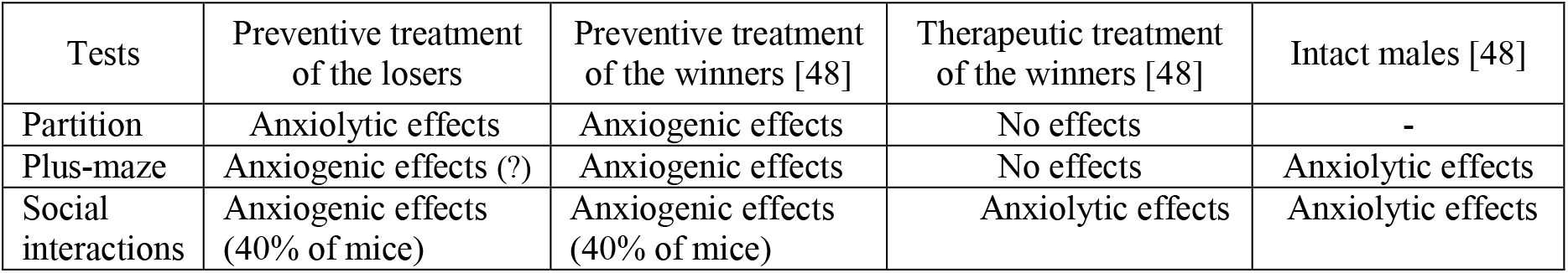
Effects of chronic LiCl treatment on the anxious behavior of mice

It was similarly shown that patients with bipolar disorder demonstrate lithium responsiveness of two types: they are good responders or poor responders [41]. Lithium is obviously a multitarget drug, which complicates understanding of its mechanism of action [59]. Lithium as a monotherapy or in combination with other drugs is effective in 60% of chronically treated patients, but response remains heterogeneous, and a large number of patients require a change in therapy after several weeks or months [60]. Therefore, there is a great need for the tools that would guide the clinicians in selecting a correct treatment strategy. Our data may be useful for understanding the variability of response to lithium in the medicine practice across individuals shown in many works [36, 37, 59]. We agree that important questions regarding the mechanism of lithium action on anxiety and depression remain open [61] despite much practice in application.

### c. Different effects of LiCl treatment in dependence on brain serotonergic activity in the mice with different social experiences in agonistic interactions

In our previous study, we discussed the development of anxiety and depression depending on the dynamic changes in brain serotonergic activity and psychoemotional states that progress in defeated mice [12]. It was shown that social defeat stress induces strong anxiety from the first days of agonistic interactions which is accompanied by the activations of the serotonergic system: the initial stage (3 days of acute social stress) is characterized by an increase of serotonin level in the hypothalamus, amygdala, and striatum. After 10 days of CSDS (stage of forming pathology) we found decreased levels of serotonin metabolite 5-hydroxyindolacetic acid in the hippocampus, amygdala and nucleus accumbens. Additionally, pharmacological desensitization and decreased number of 5-HT1A-receptors in the frontal cortex and amygdala were found. At the stage of pronounced depression (after 20 days of CSDS), there were no differences in the serotonin and its metabolite levels in all brain areas studied except hypothalamus of the losers in comparison with the control. However increased number of 5-HT_1A_-receptors in the amygdala, and decreased TPH and MAOA activities in the hippocampus have been found in the depressive mice. Later, downregulation of serotonergic genes expression in depressive mice was shown in the midbrain raphe nuclei [13, 14], which contains the majority of serotonergic neurons bodies. Taken together these data indicate the development of hypofunction of the serotonergic system at the stage of pronounced depression [12] similar with depressive patients. All these data indicated a link between duration of social defeat stress, serotonergic activity in the brain, depression and neuroplasticity disturbances that was shown also in other studies [2, 3, 7, 8]. It has been assumed that chronic preventive administration of LiCl against the background of severe stress and anxiety in animals with experience of aggression or social defeats produced an anxiogenic effect [12, 56], possibly due to activation of the serotonergic genes, and as consequences, activation of serotonergic system as side effect of LiCl.

## Conclusion

Pharmacogenomic studies have identified candidate genes that may be sensitive to antidepressants and mood stabilizers, in particular, to the lithium: as example, serotonergic *Htr2a, Htr1a, Slc6a4, Maoa, Tph* genes among numerous other genes [38]. We can assume that hyperexcitability of serotonergic neurons, induced by calcium signaling abnormalities shown earlier [62] and in our experiment (unpublished data), may be cause of overexpression of serotonergic genes under the LiCl treatment in defeated mice in our experiment. These results may be useful for understanding the mechanisms of psychotropic LiCl action through the increase of serotonergic gene expression and thereby serotonergic activity. However, it’s necessary to take into consideration that numerous genes associated both with lithium exposure and bipolar disorder were also identified [36–39, 41], and differential expression of these genes in brain tissue samples from patients and healthy controls was studied [63]: lithium exposure significantly affected 1108 genes, 702 of which were upregulated and 406 genes were downregulated. Our neurogenomic data, obtained in last years by whole transcriptomic analysis, revealed changes in the expression of mitochondrial [64], ribosomal [65, 66], monoaminergic [67–70], autistic [71] genes as well as changes in the expression of neurotrophic, transcription factors [72, 73] and collagen [74] genes specific for brain regions in mice with mixed anxiety/depression-like state. It confirms that there are many mechanisms that may account for the effects of lithium [39, 75, 76] on the neurochemical, cellular, and genomic levels. It is becoming apparent that the study of molecular mechanisms of neuroplasticity provides the most promising basis for understanding the pathogenesis of chronic anxiety, depression and anxiolytic and antidepressant efficacy. Our behavioral approach can be useful to studying effects of the lithium in psychoemotional disorders.

## Acknowledgement

We are grateful to V.N. Babenko and O.E. Redina for helpful discussion and O.A. Kharlamova for correction of manuscript text. Preparation and maintenance of experimental animals was carried out in the vivarium of the Institute of Cytology and Genetics SB RAS.

## Author Contributions

DAS received behavioral and brain data, analyzed and processed data statistically, wrote manuscript text; ILK received brain materials; AGG contributed to behavioral data acquisition, IVB organized an experiment, revised statistics and text of manuscript critically; NVT received and analyzed RT-PCR data; KOB analyzed RT PCR data; NNK performed study design, analyzed and interpreted data, wrote the main manuscript text. All authors gave final approval.

## Compliance with Ethical Standards

The authors declare that all ethical standards are met.

## Ethical Approval

All procedures were carried out in compliance with international regulations for animal experiments (Directive 2010/63/EU of the European Parliament and of the Council on the Protection of Animals Used for Scientific Purposes). The protocol for the studies was approved by Scientific Council No 9 of the Institute of Cytology and Genetics SD RAS of March, 24, 2010, N 613 (Novosibirsk).

